# Probiotic and postbiotic treatment *in situ* during a marine heat wave improves coral health and promotes specific metabolic and microbiome changes

**DOI:** 10.1101/2025.10.09.679041

**Authors:** Erika P. Santoro, Inês Raimundo, Laura Beenham, Neus Garcias-Bonet, Francisca C. García, Júnia Schultz, João Curdia, Chakkiath Paul Antony, Nathalia Delgadillo-Ordoñez, Upendra Singh, Alexandre S. Rosado, Mariusz Jaremko, Lukasz Jaremko, Raquel S. Peixoto

## Abstract

Microbial therapies are emerging as promising tools for coral protection against heat stress, yet such application was not tested *in situ*. We tested two coral-derived probiotic consortia and their heat-killed counterparts (postbiotics) on bleaching *Acropora* cf. *valida* in the Red Sea during a marine heatwave. Over 15 days, both live and one of the heat-killed treatments maintained photosynthetic efficiency (*Fv/Fm*), whereas placebo-treated corals exhibited significant thermal stress-induced decline. 16S rRNA gene sequencing profiling showed enrichment of putatively beneficial genera (e.g., *Terasakiispira spp.*, *Pseudoalteromonas spp.*), and untargeted metabolomics resolved treatment-specific metabolic signatures that differed among consortia and between live and inactivated formulations. These molecular fingerprints illuminate specific underlying protective mechanisms and host–microbiome interactions under heat stress. Together, our results position microbial therapies (probiotics and specific postbiotics) as field-validated, mechanism-informed interventions capable of minimizing impacts on corals during real-world thermal extremes and provide design cues for scalable deployment.

## Introduction

Marine ecosystems are facing unprecedented global decline, driven by a combination of local and global stressors^1–4^. Recent mass bleaching and mortality events highlight the urgent need for proactive, science-based strategies to promote coral reef assisted restoration focused on strengthening coral reef resilience^5–7^. Coral holobionts depend heavily on microbial interactions for their health and survival, in^1–4^ particular, bacteria play a critical role in enhancing stress tolerance and facilitating adaptation to changing environmental conditions^8–12^. However, environmental stressors can disrupt these microbial communities, often triggering shifts toward more pathogenic or dysbiotic states, which not only compromise coral health but can also impact the broader reef ecosystem^13,14^. The conservation and rehabilitation of corals and their microbial diversity with the refinement of active biotechnological interventions is one of the climate priorities for this decade^15–17^.

Microbial therapies can restore or rehabilitate native microbiomes following a disruption caused by different environmental impacts (e.g., pathogen infection, antibiotic treatment, changes in temperature and pollution)^18–21^ and have been highlighted as a biotechnological conservation strategy to reduce biodiversity loss^22^. The use of probiotics, for example, can restructure the microbiome and promote the health and growth of corals *ex-situ* ^22^, which are threatened with functional extinction due to local and global impacts, such as marine heatwaves, by 2030^4,23^. Probiotics are typically composed of single or combined consortia^24,25^ of living microbes that, when applied in the right concentration and regime, can enhance an organisms’ health^26^. They can be obtained from different sources but seem to be more effective when isolated from healthy individuals of the target species and selected based on specific beneficial traits^24,27^. Probiotics obtained from corals and other marine organisms, called Beneficial Microorganisms for Corals (BMC)^28,29^, boosted coral growth or minimized the effects of different types of impacts, such as thermal stress, pathogen infection, and oil spills in laboratory trials ^30–39^.

In addition to probiotics, alternative approaches to microbial therapy have been suggested to support coral restoration and rehabilitation efforts, such as microbial transplantations^10^ and postbiotics^24^, which are heat-killed probiotic microorganisms and/or their bioactive components^26^. These inactivated cells or their components have been reported as beneficial to humans and plants through the activation of immune and/or other responses in the host^40–48^. Whether postbiotics trigger microbiome restructuring and/or host responses in humans and plants is usually determined by their compositional lipopolysaccharides, lipoteichoic acids, peptidoglycans, and exopolysaccharides^40,49–51^, which is why dead cells cannot be used as negative controls in studies of probiotics^24^. Establishing whether postbiotics can match the effects of probiotics in corals is crucial, given their potential advantages in shelf-life, safety, cost, and efficacy ^24,40,42,52^.

Another critical knowledge gap regarding the use of different microbial therapies in corals is whether these treatments will be feasible for real-world applications, including to counteract thermal stress, and what are the underlying mechanisms triggered by these treatments^53,54^. Although data indicate that the coral microbiome can be effectively restructured through the use of probiotics *in situ*^55^, this field experiment was conducted under normal conditions, where all corals were healthy and not exposed to thermal stress.

Here, we report a field study using probiotic and postbiotic therapies during a marine heatwave. By (i) comparing for the first time the use of viable and non-viable combinations of the same BMC (i.e., the same assemblage of coral probiotics and postbiotics); (ii), testing the application of these microbial therapies *in situ*, and (iii), exploring microbiome and metabolic changes promoted by each of these treatments, and how these consortia correlate with specific metabolic signatures and health improvements in heat-stressed corals we gain substantial insight about the mechanisms underlying microbial therapies. We therefore demonstrate the viability of both probiotic and postbiotic interventions as a tool for coral reef restoration during thermal stress.

## Results

### Probiotic consortia improves coral health during heat stress while only a specific corresponding postbiotic consortium produced similar effect

Two probiotic consortia (BMC1 - two *Pseudoalteromonas galatheae*, two *Cobetia sp.*, one *Halomonas sp.* and one *Sutcliffiella sp.;* BMC2 - one *Halomonas piezotolerans*, one *Pseudoalteromonas sp.*, one *Pseudoalteromonas lipolytica* and one *Cobetia sp.*) both containing bacterial strains isolated from Red Sea corals and selected by their beneficial traits, as well as their corresponding postbiotic counterparts (D_BMC1 and D_BMC2) were used for this study. During a 15 day field experiment, placebo or one of the consortia were applied to 25 (n = 5 per treatment group) heat-stressed *Acropora cf. valida* colonies that were showing signs of bleaching in the central Red Sea during a marine heat wave. During this period, the daily average seawater temperature ranged from 30.87 °C to 31.95 °C, resulting in a relative Degree Heat Week (rDHW) of 12.4 °C-weeks at our site based on daily mean in situ measurements ^56^ - Table S1). The effect of the probiotic and postbiotic consortia on coral health status was measured in two ways: qualitative bleaching scores (color monitoring) and endosymbiotic algae photosynthetic efficiency. Color monitoring was based on underwater pictures taken of the colonies using the coral health chart as a color reference. At the beginning of the experiment (T0), most of the colonies were pale due to the heat stress, with their colors being, on average, equivalent to the category D4 of the coral health chart. Colonies treated with Placebo, BMC1, BMC2 and D_BMC2 showed a slight improvement in their bleaching score after 15 days of experiment (T1), with an average score of D5 on the coral health chart; D_BMC1 treatment did not alter color scoring (Fig. S1).

The photosynthetic efficiency (*Fv/Fm*) of the endosymbiotic algae on treated corals was measured using imaging-pulsed amplitude modulation (PAM). Placebo-treated corals exhibited a significant decrease in *Fv/Fm* from T0 to T1 (emmeans post hoc test: p = 0.0473 - Fig. 1A, Table S2). Additionally, corals treated with D_BMC1 showed a significant *Fv/Fm* decline in T1 compared with T0 (emmeans post hoc test: p= 0.0136, Table S2). Conversely, corals treated with BMC1, BMC2 and D_BMC2 did not show significant differences in *Fv/Fm* between T0 and T1 (emmeans post hoc tests: p= 0.9845, p= 0.5783, and p= 0.2078, respectively; Table S2; Fig. 1A, Table S2), suggesting that these treatments induced maintenance of photosynthetic efficiency during the marine heatwave. Moreover, comparisons of *Fv/Fm* values between placebo and each treatment group at T1 revealed that corals treated with BMC1 and D_BMC2 exhibited significantly higher *Fv/Fm* than placebo-treated corals (ANOVA - F = 7.106, p= 0.028; ANOVA - F = 6.653, p= 0.033, respectively). No significant differences were detected between D_BMC1 and placebo treatments (ANOVA - F = 0.264, p= 0.64 - Fig. 1B, Table S2). Therefore, corals treated with BMC1 and D_BMC2 improved significantly the health of *Acropora cf. valida* within 15 days, as evidenced by decreases in bleaching scores and improvements in photosynthetic efficiency.

**Figure 1:**
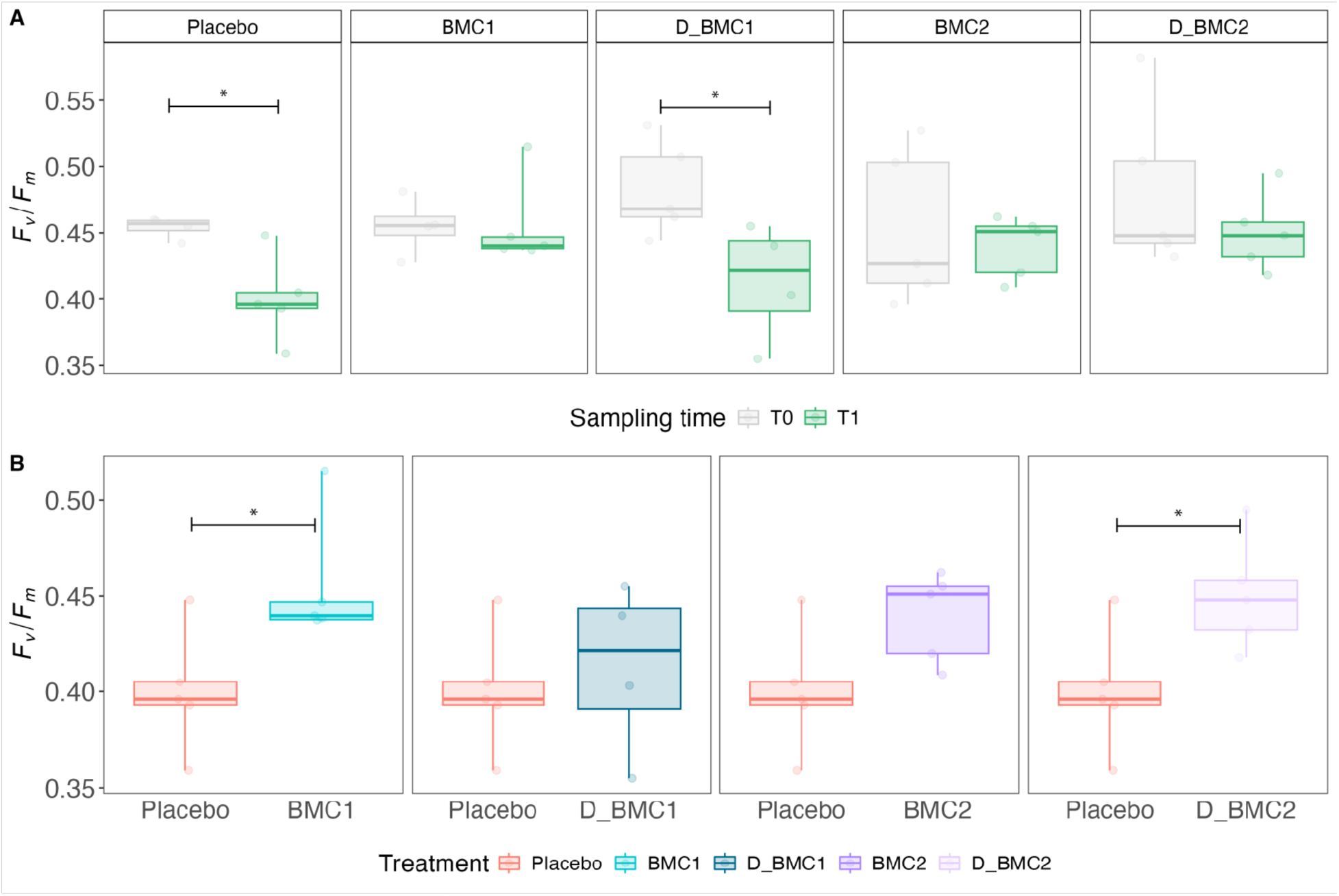
Differences in photosynthetic efficiency between corals treated with probiotics, postbiotics or Placebo. (**A**) Photosynthetic efficiency (*F_v_/F_m_*) at T0 and T1 for the different treatment groups. (**B**) Pairwise comparison of *F_v_/F_m_* between all treatments and placebo at sampling time T1. The boxplots show the median (center line) and the first and third quartiles (lower and upper bounds). Biological replicates per sampling time and treatment (n=5).

### Bacterial strains composing probiotic and postbiotic consortia are enriched in treated corals

To test whether strains from consortia 1 and 2 were enriched in the microbiomes of treated corals, we used the full-length 16S rRNA gene sequence of each strain to search for matches among the amplicon sequence variants (ASVs). ASVs with ≥99% identity with the consortium-one strains (*Cobetia* spp., *Halomonas* sp., *Pseudoalteromonas galathea*e and *Sutcliffiella* sp.) and consortium-two strains (*Cobetia* sp*., Halomonas piezotolerans* sp., and *Pseudoalteromonas* sp.) were found in placebo, probiotics-(BMC1 and BMC2) and postbiotics-treated (D_BMC1 and D_BMC2) samples (Fig. 2). Importantly, no ASVs with ≥99% sequence identity to *Pseudoalteromonas lipolytica* were detected in the dataset, and therefore this strain was not included in the plots of this analysis. ASVs with ≥99% identity with *Cobetia spp.* were significantly different at T1 among treatments (Kruskal Wallis - H = 6.26, p= 0.043 for *Cobetia sp.* in corals treated with the consortium-one either dead or live; Kruskal Wallis - H = 8.34 p = 0.0155 Cobetia *sp.* in corals treated with the consortium-two either dead or live - Table S3). Specifically, D_BMC1 treated corals showed a higher relative abundance of *Cobetia spp.* ASVs at T1 than placebo treated corals (Dunn test - p*adj* = 0.019 - Fig. 2, Table S3). Similarly, corals treated with BMC2 (Dunn test - p*adj* = 0.025) and D_BMC2 (Dunn test - p*adj* = 0.008) were also enriched with *Cobetia sp.* compared with placebo at T1 (Fig. 2, Table S3).

**Figure 2:**
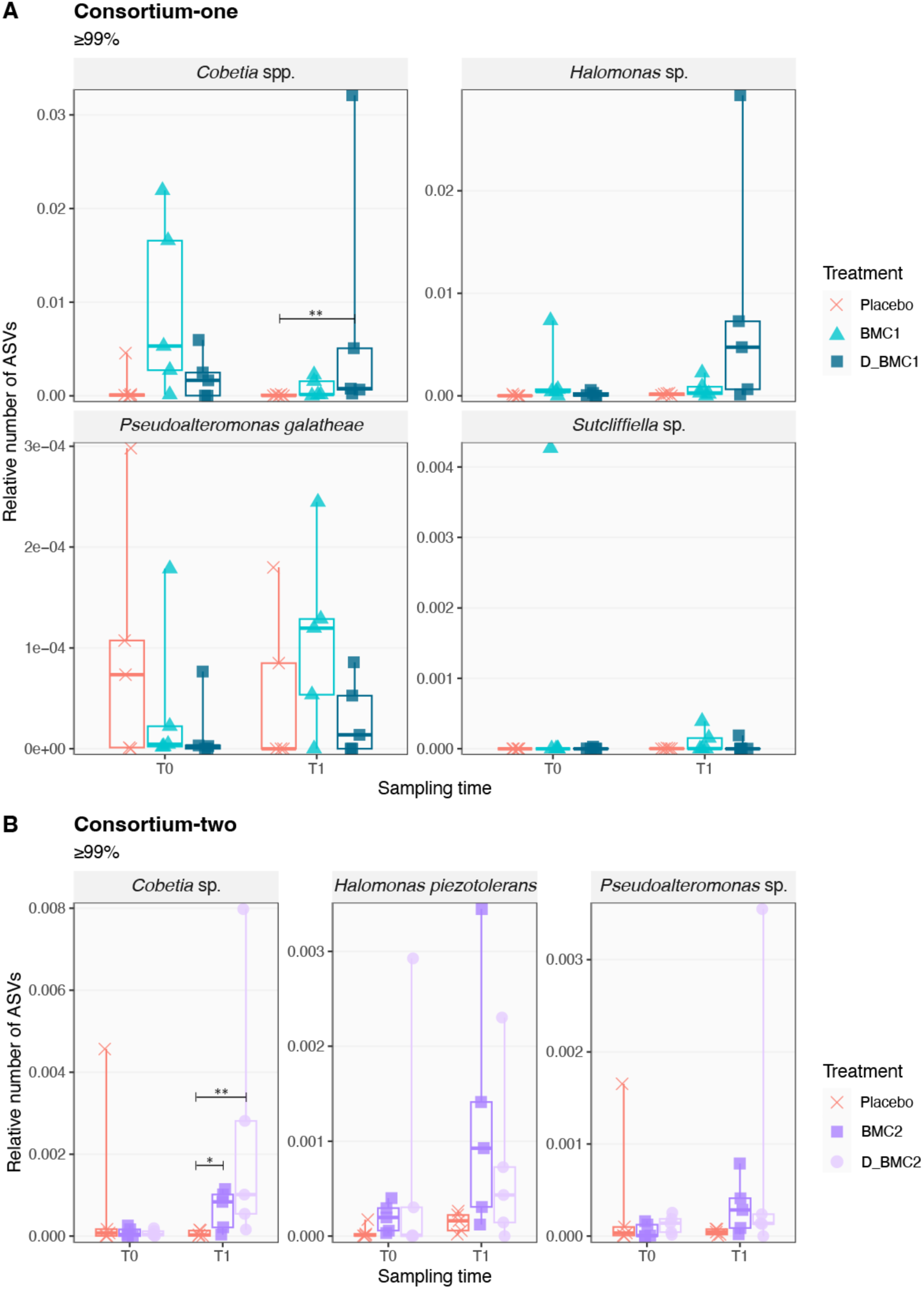
Enrichment of bacterial strains from consortia-one and -two in treated corals. The graphs show the relative numbers of ASVs with ≥99% identity with the indicated bacterial strains. The depicted boxplots show the median (center line) and the first and third quartiles (lower and upper bounds). (**A**) Treatment placebo, BMC1 and D_BMC1. (**B**) Treatment with placebo, BMC2 and D_BMC2. Pairwise significance determined by Dunn’s test is indicated in the graphs as follows: p*adj* ≤ 0.05(*), p*adj* ≤ 0.01 (**), p*adj* ≤ 0.001 (***).

### Each microbial therapy triggers the enrichment of specific ASVs

We further assessed the effects of the treatments with probiotic and postbiotic consortia in the coral microbiome through the analysis of 16*S* rRNA gene sequencing of coral samples taken at T0 and T1. After removing potential kit contaminants as well as ASVs identified as mitochondria or chloroplasts and singleton sequences, a total of 10,688 ASVs were retained for downstream analysis. Read-rarefaction was applied only on alpha-diversity analysis (see Methods), after which a total of 5,157 ASVs were retained. Even though probiotic and postbiotic treatments induced differential responses on coral photosynthetic efficiency (*F_v_/F^m^* - Fig. 1), no overall changes were observed in alpha (Fig. S2, Table S4) or beta-diversity (Fig. S3A, Table S5) between treatments or sampling time.

In all treatment groups, the coral microbiome was dominated by Endozoicomonadaceae in both T0 (58.9 % in Placebo, 55.5 % in BMC1, 36.7 % in D_BMC1, 73.8 % in BMC2, 57.1 % in D_BMC2 - Fig. S3B) and T1 (44.8 % in Placebo, 59.3 % in BMC1, 45.8 % in D_BMC1, 42.1 % in BMC2, 39.5 % in D_BMC2 - Fig. S3B, Table S6). Since such strong dominance may mask smaller compositional changes, we assessed differentially abundant (DA) taxa between placebo- and probiotic- or postbiotic-treated samples at T1. DESeq2 analysis revealed a high number of DA ASVs between treatments, and only ASVs demonstrating significant enrichment (FDR adjusted p <0.01) are reported (Fig. 3, Table S7).

**Figure 3.**
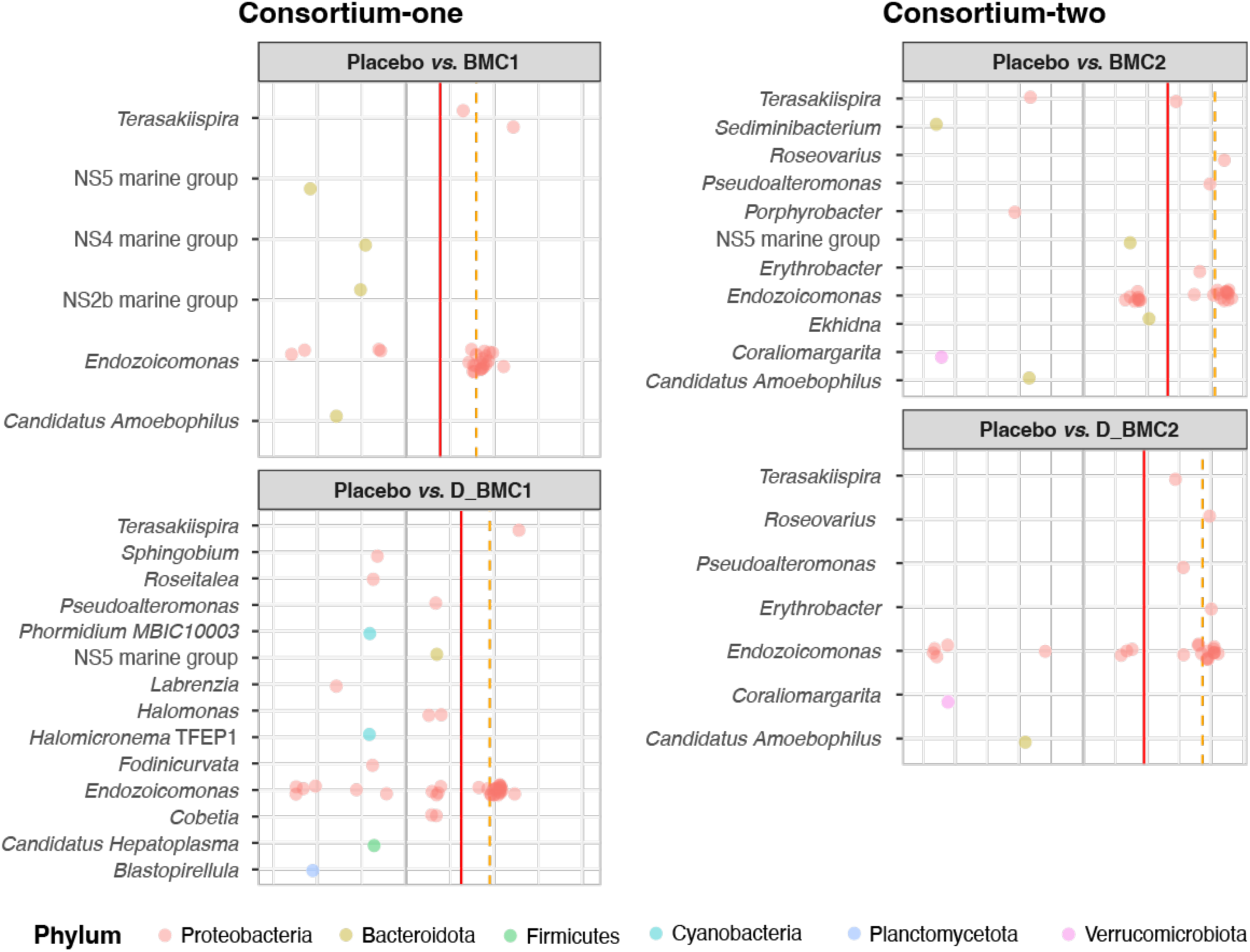
Significant differential abundance of ASVs according to consortia treatment. Only ASVs present in three or more samples per treatment group (n = 5) were included in the analysis. Each point represents a single ASV that increased or decreased significantly (FDR corrected p < 0.01) at T1 compared with placebo and comparing probiotic to postbiotic. Each row and dot color corresponds to a bacterial genus. The red colored line within each box represents the mean log2FoldChange for ASVs with significant changes. The dashed orange colored line represents the median log2FoldChange for ASVs with significant changes. The gray colored line at 0 log2FoldChange indicates no effect or no interaction.

Interestingly, coral treated with either probiotic or postbiotic were enriched almost exclusively with ASVs belonging to Pseudomonadota when compared to placebo, which includes *Endozoicomonas*, *Halomonas*, *Terasakiispira*, *Roseovarius*, *Pseudoalteromonas* and *Erythrobacter* (Fig. 3, Table S7). Conversely, placebo-treated corals were enriched with ASVs from Bacillota, Bacteroidota, Actinobacteriota, Cyanobacteria, Verrumicrobiota and Planctomycetota (Fig.3, Table S7). ASVs identified as *Endozoicomonas* were enriched more prominently in samples treated with either probiotics or postbiotics in comparison with placebo (BMC1 - 14 DA ASVs, LFC values from 16.842 to 22.125, FDR padj< 0.01; BMC2 - 17 DA ASVs, LFC values from 6.570 to 23.213, FDR padj< 0.01; D_BMC1 - 22 DA ASVs, LFC values from 16.510 to 24.624, FDR padj< 0.01; D_BMC2 - 15 DA ASVs, LFC values from, FDR padj< 0.01 - Fig. 3, Table S7). ASVs belonging to *Terasakiispira* genus were also enriched only in corals treated with probiotics and postbiotics but not in the placebo-treated corals (BMC1 - LFC = 12.96 and 24.32, FDR adjusted p< 0.01; D_BMC1 - LFC = 25.587, FDR adjusted p< 0.01; BMC2 – LFC =14.513, FDR adjusted p< 0.01; D_BMC2 - LFC = 14.408, FDR adjusted p< 0.01; Fig. 3, Table S7). ASVs identified as *Pseudoalteromonas* were also enriched in treated corals (except BMC1) in comparison with placebo (D_BMC1 - LFC = 6.765, FDR adjusted p< 0.01; BMC2 - LFC = 19.799, FDR adjusted p< 0.01; D_BMC2 - LFC = 15.639, FDR adjusted p< 0.01; Fig. 3 Table S7). Exclusively corals treated with BMC2 or D_BMC2 were significantly enriched with ASVs belonging to *Roseovarius* genus when compared to placebo (BMC2 - LFC = 22.052, FDR adjusted p< 0.01; D_BMC2 - LFC = 19.747, FDR adjusted p< 0.01). Additionally, treatment with both BMC2 and D_BMC2 induced an enrichment of ASVs belonging to the same genera, such as *Terasakiispira*, *Roseovarius, Pseudoalteromonas*, *Erythrobacter* (LFC = 18.209 and 20.014, FDR adjusted p <0.01, BMC2 and D_BMC2 respectively) and *Endozoicomonas* (Fig.3, Table S7). These findings show that both probiotic and postbiotic treatments enriched shared bacterial genera, though some taxa responded uniquely to specific treatments.

### Treatment signature: In-situ short-term treatments using probiotics alter coral metabolic profiles

To assess how probiotic and postbiotic treatments affect the metabolic profiles of the coral holobiont, potentially supporting adaptation to heat stress, we used NMR to analyze secondary metabolites in coral samples at T0 and T1. A total of 21 distinct metabolites, as well as 2 unknown metabolites, were identified across treatments. Initial analysis revealed a strong overlap in ^1^H NMR spectra between all groups at T0, before the use of microbial treatments, indicating the metabolic profiles were similar amongst all corals tested at the beginning of the study (Fig. S4, Table S8).

At T1, the metabolomic profiles of corals differed significantly among treatments (PERMANOVA - F = 3.562, R² = 0.428, p = 0.001; Fig. 4A, Table S8). BMC1-treated corals had the most divergent metabolomic profile, differing significantly from every other treatment (see Table S8 for pairwise PERMONOVA). Corals treated with D_BMC2 also exhibited significantly different metabolomic profiles compared to those treated with BMC1 (pairwise PERMANOVA: F = 4.128, R² = 0.340, p*adj* = 0.022), and D_BMC1 (F = 2.491, R² = 0.237, p*adj* =0.022), but not from BMC2 or placebo-treated corals (Table S8). These differences were primarily driven by three metabolites: Formate (ANOVA - F = 6.722, R²= 0.585, p*adj* = 0.027), Trimethylamine N-oxide (ANOVA - F = 6.124, R²= 0.563, p*adj* = 0.027) and Isoleucine (Kruskall-Wallis - H = 14.317, ε^2^= 0.543, p*adj* = 0.048 - Fig 4B, Table S8). Formate and Trimethylamine N-oxide seem to have significantly higher concentrations in BMC1 compared to Placebo (Tukey, p*adj* = 0.0008; Dunn test p*adj* = 0.002, for Formate and Trimethylamine N-oxide respectively) and to D_BMC2 (Tukey, p*adj* = 0.010; Dunn test p*adj* = 0.006, for Formate and Trimethylamine N-oxide respectively). Trimethylamine N-oxide also demonstrated to be significantly higher in BMC1 when compared to D_BMC1 (Tukey, p*adj* = 0.040). Additionally, Isoleucine appeared to have significantly higher concentrations in D_BMC1 when compared to Placebo (Dunn test, *p*adj = 0.020) and D_BMC2 (Dunn test, *p*adj = 0.020 - Fig. 4C, Table S8).

**Figure 4.**
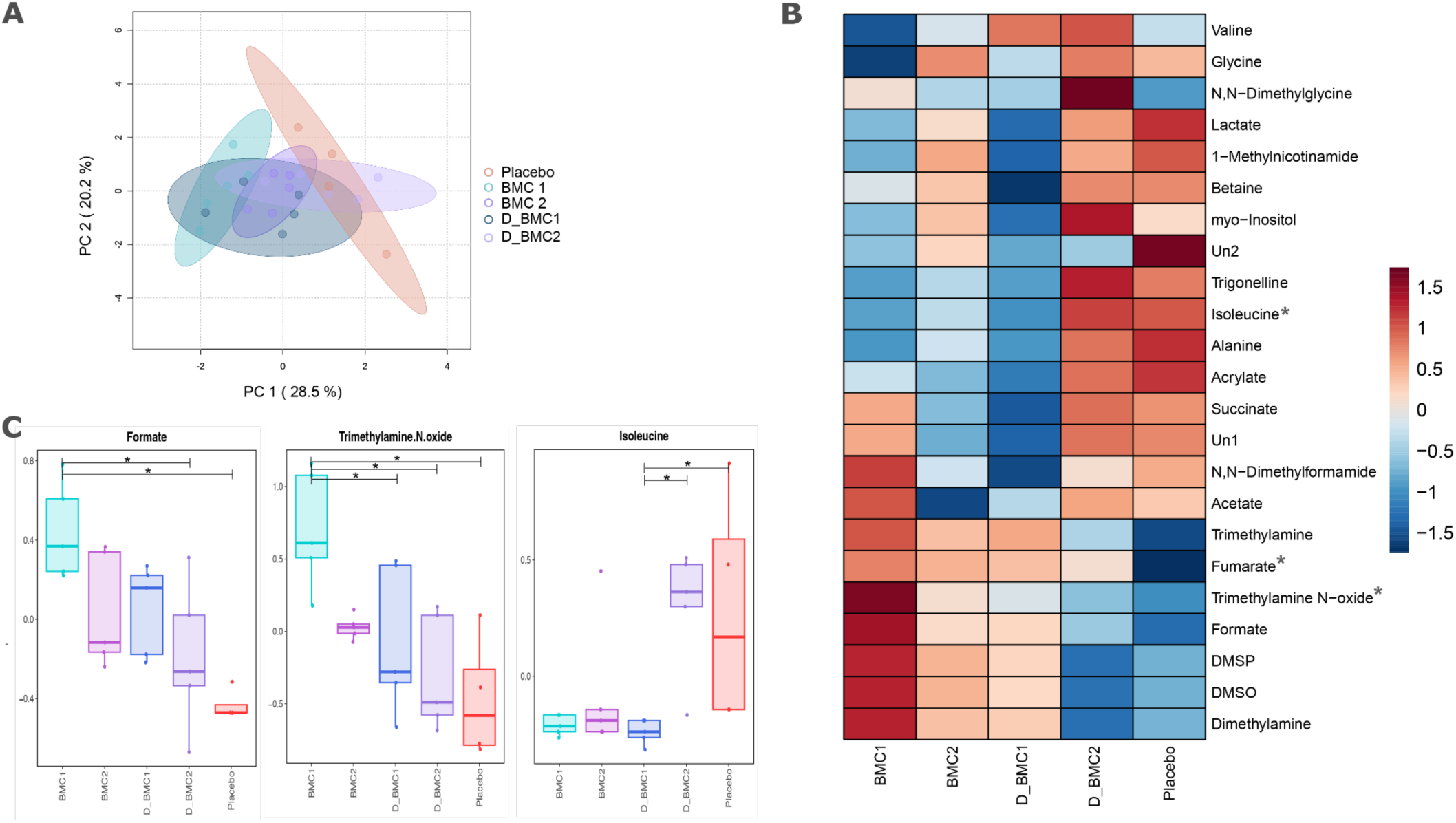
Probiotic and postbiotic treatments differentially alter coral metabolic profiles: (**A**) Principal Components Analysis (PCA) of the ^1^H NMR dataset comparing the metabolic patterns of coral samples from each of the treatments at T1. (**B**) Heatmap of metabolite dataset by treatment, stars (*) identify the metabolites with significant differences by treatment in T1 (one way ANOVA or Kruskal-Wallis, p≤ 0.05). (**C**) Boxplot of significantly different metabolite concentrations per treatment at T1, where significant pairwise comparisons were determined by Tukey or Dunn’s posthoc, and is identified by stars, as follows: p*adj* ≤ 0.05(*), p*adj* ≤ 0.01 (**), p*adj* ≤ 0.001 (***).

Correlation analysis between bacterial genera composing consortia-one and -two and the metabolites detected through ^1^H NMR at T1 revealed differential metabolic profiles according to treatment (Fig. 5). For example, *Cobetia spp.* showed positive correlations with X1 Methylnicotinamine (D_BMC1 r = 0.87 p ≤ 0.05) and an unknown metabolite (D_BMC2 r = 0.90, p ≤ 0.05) in corals treated with postbiotics. Similarly, *Halomonas* demonstrated positive correlations with several metabolites, including NN Dimethylglycine (BMC1 r = 1.00, p ≤ 0.05), Alanine, Betaine, and Glycine (BMC2 r = 0.70, 0.90, and 0.90, respectively, p ≤ 0.05), as well as X1 Methylnicotinamine, Trimethylamine, and Valine (D_BMC2 r = 0.90, 0.90 and 0.90 respectively, p≤ 0.05). *Pseudoalteromonas* correlated with Trimethylamine N-oxide and myo-Inositol (BMC1 r =1.00 and 1.00, respectively, p≤ 0.05), as well as Dimethylamine, Formate, DMSO, DMSP, and Trigonelline (BMC2 r = 1.00, 0.70, 1.00, 1.00 and 0.90, respectively, p≤ 0.05). Interestingly, probiotics and postbiotics treatments seem to enhance correlations between beneficial members and the detected metabolites when compared to placebo. For example, in placebo-treated corals only *Pseudoalteromonas* showed positive correlations (unknown metabolite, Glycine, Trimethylamine, and Trimethylamine N-oxide (r = 1.00, 1.00, 1.00 and 1.00, respectively, p≤ 0.05)), while more than one beneficial member (*Cobetia spp., Halomonas spp. and/or Pseudoalteromonas spp.*) correlated with metabolites when treated with either probiotics or postbiotics (Fig. 5). This gives us insights on how probiotics and postbiotics treatments interact with the metabolism of coral holobionts, also opening the pathway for potential markers for heat-stress.

**Figure 5.**
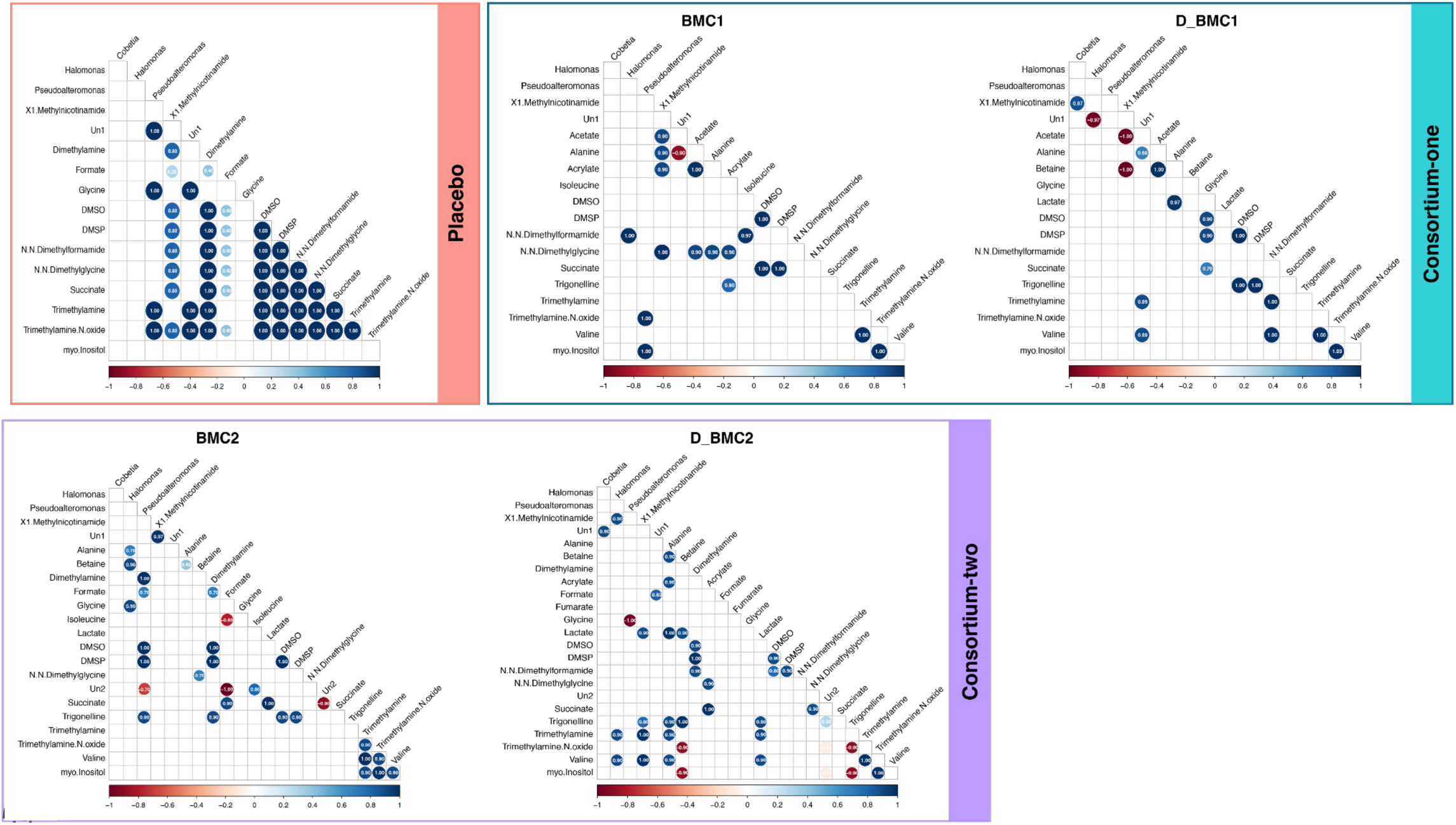
Genera from consortia-one and two exhibit significant correlations with metabolites across probiotic, postbiotic, and placebo treatments at T1. Correlation matrices showing co-occurrence of the consortia genera and the metabolites found to be significantly detected within probiotics (BMC1 and BMC2), postbiotics (D_BMC1 and D_BMC2) and Placebo at T1. The interactions are color-coded based on their linear positive (blue) or negative (red) correlations, of which range between -1 and 1. The color depth indicates the strength of the correlation. Coefficient of correlation is reported.

## Discussion

Probiotics have been shown to protect corals from thermal stress in controlled settings^8,38,39^, prevent mortality in mesocosm experiments^32^, and restructure coral microbiomes *in situ* without detectable environmental side effects^55^. They have also benefited other reef organisms such as fish^57,58^ and sponges^57,58^. However, the benefits of using probiotics to improve coral health during heat stress *in situ* and the use of postbiotics (*in situ* or *ex situ*) for corals have never been reported. Here, we demonstrate that i) different microbial therapies, including postbiotics (dead cells), can improve coral health during a marine heatwave *in situ*, and ii) each consortium, as well as their status (either dead or alive) induces distinct physiological, microbial and metabolic responses. More specifically, we show that while both probiotic consortia promoted health improvements, only one postbiotic assemblage (D_BMC1) was effective. iii) We also identify specific changes in the coral microbiome and metabolic profile that are linked to the mechanisms underlying the observed health benefits.

The field experiment took place in the central-eastern Red Sea during a marine heat wave in summer 2022. We monitored *Acropora* cf. *valida* corals showing signs of heat stress (as reflected by their pale coloration) treated *in situ* with different formulations of probiotics (BMC1 and BMC2) and their corresponding postbiotics (D_BMC1 and D_BMC2), for a total of 15 days. Colonies treated with placebo had a significant drop in their *Fv/Fm* values from T0 to T1, indicating temperature-related impairment of photosystem II electron transport of the associated Symbiodiniaceae, as expected for corals under heat stress^59^. By contrast, colonies treated with probiotics (BMC1 and BMC2) and the D_BMC2 postbiotic consortia maintained photosynthetic efficiency when comparing T0 with T1.

The protective effect of probiotics in helping corals withstanding heat stress and evading mortality has been previously demonstrated in mesocosm experiments^8,38,39,60^; however, this study marks the first *in-situ* validation of their efficacy, reinforcing the use of microbial therapies as a tool for coral restoration in “real world scenarios”. Notably, the postbiotic consortium D_BMC2 replicated the beneficial effect of its live probiotic counterpart on coral photophysiology (*Fv/Fm*) under thermal stress, although apparently through different mechanisms, as indicated by the different metabolic outcomes observed. In other systems, such as humans and plants, inactive cells and their components have been shown to confer health benefits by modulating immune responses and other host physiological processes^40–48^. However, the full mechanisms by which these inactivated cells or their bioactive metabolites can exert such effects in coral remains to be further investigated and is of great interest., the observed decline in *Fv/Fm* from T0 to T1 in corals treated with postbiotic consortium-one (D_BMC1) highlights the specificity of host responses to consortia composition and supports the notion that dead cells cannot be assumed as neutral or inert controls in probiotic studies^24,40,42,52^.

Additional to health improvements, we observed a significant increase (sequences with ≥99% identity with the strains used in each consortia - see Fig 4.) of *Cobetia spp.* in postbiotics-treated corals when compared with placebo at T1, suggesting the enrichment of some of the BMC strains into the microbiome of the treated corals. The application of probiotics *in situ* was recently tested in healthy corals, and resulted in the incorporation of two BMC genera in probiotic-treated corals only after a long period of time (two months)^55^. Our results show that consortia enrichment is faster under stress conditions, occurring within 15 days, and it does correlate with physiological improvements *in situ* depending on the consortia. Regardless of its correlation with health improvements (D_BMC2) or not (D_BMC1), it’s worth it to highlight the enrichment of BMC strains and other beneficial members of the coral microbiome in postbiotic-treated corals. This indicates that dead cells can also trigger microbiome restructuring, possibly by metabolic favoring, as observed in other hosts^24,42^, and in this case specifically favoring live *Cobetia spp.* counterparts within the coral microbiome.

Even though treatment with probiotics (BMC1 and BMC2) and postbiotic (D_BMC2) induced changes in coral photophysiology, microbial diversity and composition remained stable, likely due to the dominance of Endozoicomonadaceae across all treatments and sampling time. Members of the Endozoicomonadaceae family are the most prevalent and consistently found bacteria associated within Cnidaria^11,55,61–63^ and their ubiquitous presence may mask smaller (but yet important) shifts. We further used a higher-resolution taxonomic analysis (DESeq) to i) investigate the improvements in *Fv/Fm* and incorporation of the BMC-*Cobetia* spp. were followed by shifts in the microbiome, and ii) whether these treatments could modulate the coral microbiome, despite its association with coral health improvements. Treated corals were enriched with *Pseudoalteromonas* spp. and *Terasakiipira* spp. compared with placebo. *Pseudoalteromonas* is a well-known coral tissue-associated genus that has been validated as a coral probiotic^31,34,38^. Although *Terasakiipira* is a relatively novel and understudied genus within the Halomonadaceae family (Zepeda et al. 2015), our findings highlight its potential relevance. Its close phylogenetic relatedness with *Halomonas*, a well-known coral-tissue associated bacterial genus that has been validated as a coral probiotic^8,38,34^, suggests *Terasakiipira* may be a promising candidate for further investigation as a beneficial coral symbiont genus. Some bacteria taxa were more selectively enriched depending on the consortia applied. For example, ASVs identified as *Roseovarius spp.* and *Erythrobacter spp.* were exclusively enriched in consortia-two treated corals (either probiotic or postbiotic), but not in consortia-one treated corals. *Roseovarius spp.* has been suggested as beneficial for corals due to its potential to produce antioxidants (such as DMSP), participate in the nitrogen and carbon cycle, and also due to its ROS-scavenging ability^64^). *Erythrobacter*, a common bacteria found in the surface mucus layer of Red Sea corals^65^, is an aerobic anoxygenic phototrophic bacteria that is hypothesized to use light as an additional energy source, similar to symbiotic algae^66,67^. This ability may reduce the consumption of organic carbon, potentially easing metabolic demands on the coral holobiont during stress^68^. Importantly, placebo-treated corals showed enrichment of ASVs belonging to more diverse phyla (Bacillota, Bacteroidota, Actinobacteriota, Cyanobacteria, Verrumicrobiota and Planctomycetota) while probiotic and postbiotic-treated corals were mostly enriched with Pseudomonadota, which aligns with the principle of which hosts microbiome appear to be less ordered when stressed^11,69,70^.

Interestly, in some cases, significant changes in the metabolic profiles followed the same trend of the health status. For example, the metabolic profile of BMC1-treated corals showed significant divergences from corals treated with placebo. More specifically, trimethylamine N-oxide and Formate was significantly enriched in BMC1 in comparison with placebo-treated at T1. Trimethylamine N-oxide is a known osmolyte compound that maintains holobiont homeostasis, and was found to be down-regulated in corals exposed to heat stress^71,72^. Formate, a key metabolite in one-carbon metabolism, contributes to nucleotide synthesis and DNA methylation via S-adenosylmethionine (SAM)^73^. Given that DNA methylation changes have been linked to coral stress responses^74,75^, elevated formate levels could be associated with enhanced phenotypic plasticity and thermal resilience. Epigenetic modifications allows organisms to rapidly adjust to environmental stress, offering a faster and flexible mechanism of acclimatization compared to genetic adaptation^74,76,77^. Moreover, probiotics were recently suggested and demonstrated to promote such epigenetic changes, the use of postbiotic may also be correlated with coral health improvements via epigenetics changes^78^.

The treatment with probiotic and/or postbiotic also seems to determine the correlation between BMC genera with specific metabolites. *Pseudoalteromonas spp.*, for example, was the only BMC strain to correlate with metabolites in placebo-treated corals. However, the presence of *Pseudoalteromonas spp.* not only correlated with trimethylamine N-oxide, but also with myo-inositol in BMC1-treated corals, and with dimethylsulfoxide (DMSO), dimethylsulfoniopropionate (DMSP), and trigonelline in BMC2 corals. DMSP and myo-inositol are important osmoregulators^79–83^ usually produced by dinoflagellates for stress protection ^83,84^ and hypothesized to mediate signaling with the coral host^84,85^. While certain marine bacteria such as *Vibrio coralliilyticus* seem to use myo-inositol as a sole carbon source to enhance growth during their growth lag-phase^86^, this ability remains uncharacterized in *Pseudoalteromonas spp.*. However, if *Pseudoalteromonas spp.* can similarly utilize myo-inositol, it may have a competitive advantage in colonizing coral, which is particularly noteworthy when considering microbial treatments. Additionally, DMSP can buffer reactive oxygen species^87^ and its degradation generates antimicrobial compounds (DMSO) that may also help control pathogens^88–90^. This buffering and protective mechanism of DMSP has been shown to be a beneficial mechanism for coral survival after probiotic inoculations under heat stress^32^. To further elucidate the mechanisms driving beneficial metabolic shifts that elicit thermal tolerance during microbial therapies, integrative approaches involving transcriptomic and epigenetic analyses are needed.

Overall, we show that the inoculation of coral colonies with probiotics (BMC1 and BMC2) and a specific consortia of postbiotics (D_BMC2, instigate rapid and specific changes in the coral-associated microbiome and their metabolomic and physiological traits. Our findings also indicate that probiotic enrichment is effective under thermal stress conditions and should be considered in future microbial therapy interventions to protect corals during marine heatwaves. Our results demonstrate that microbial therapies can promote coral health improvements in real-world trials but this is dependent on the (type of) consortia. While both probiotic consortia (BMC1 and BMC2) improved coral health parameters, only one postbiotic consortium (D_BMC2) mirrored the effects of its live counterpart, highlighting the specificity of host response and the notion that dead cells cannot be used as treatment-controls. These results underscore the importance of tailored microbial therapies as a proactive strategy for coral conservation and restoration under climate change. Future work should focus on optimizing strain and/or metabolites selection and delivery methods to develop reliable, scalable interventions for protecting reefs during marine heatwaves.

## Materials and methods

### Study site and permits

This study was conducted at the “Red Sea Research Center Coral Probiotic Village” (CPV - 22° 18.3071’ N, 38° 57.8741’E), a mid-shore reef in the central-eastern Red Sea, North Al Fahal, Thuwal, Saudi Arabia (Fig. 6A). The CPV consists of a shallow sheltered area spanning about 500 m^2^, with maximum depth ranging from 8-10 m, about 15 km off-shore from King Abdullah University of Science and Technology (KAUST). The CPV is a multidisciplinary underwater laboratory established to test the efficacy of probiotics and other restoration strategies *in situ*. The area has been thoroughly characterized by Garcias-Bonet and colleagues (^56^ and its biotic and abiotic parameters are constantly monitored. All the fieldwork and sampling procedures were conducted in compliance with 22IBEC003 permits and KAUST’s regulations.

**Figure 6:**
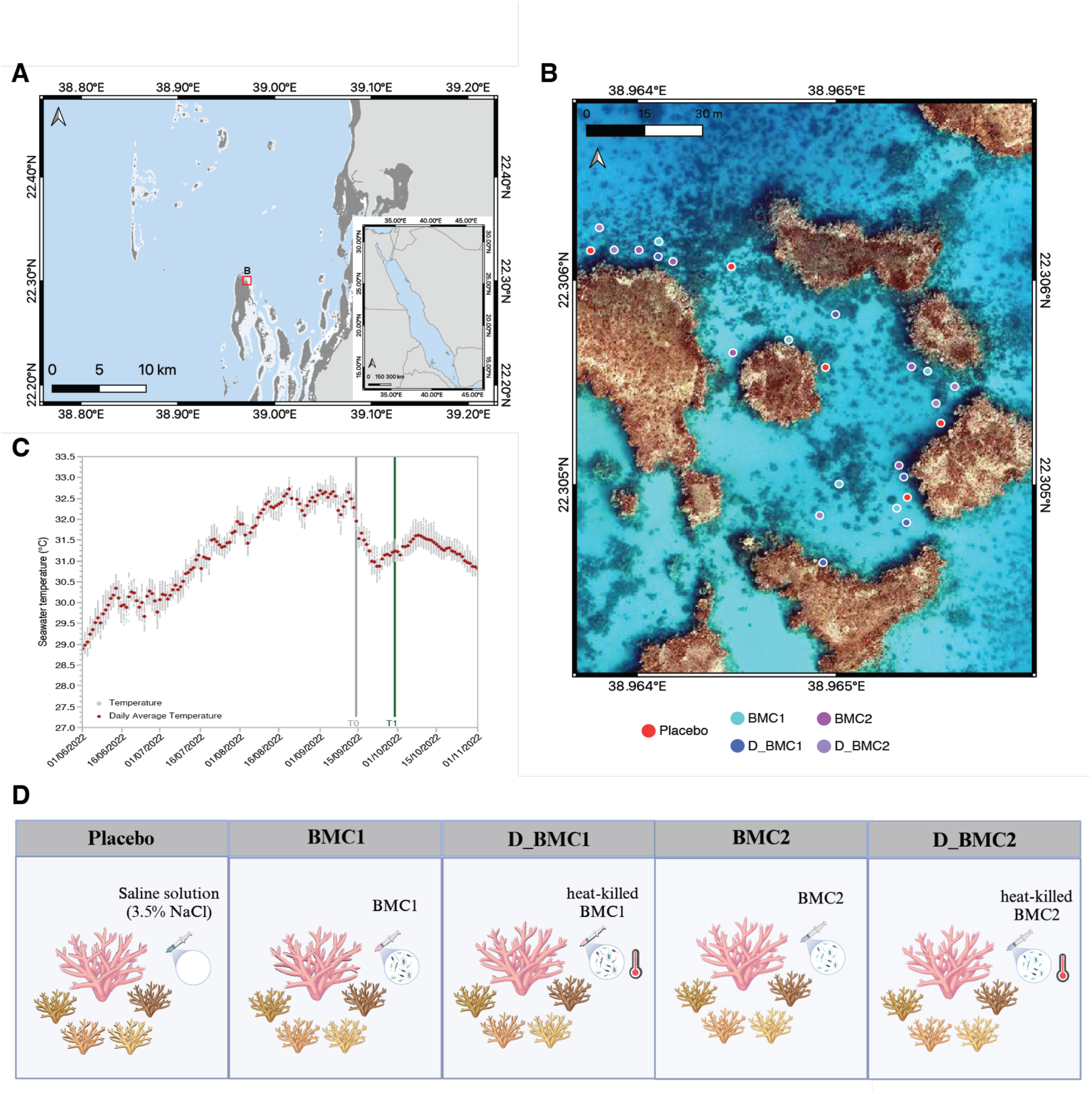
Study area and experimental design. (**A**) Study site in the Central Red Sea. (**B**) Drone image of the study site ^56^ and coral colonies assigned per treatment, where Placebo is red, BMC1 aqua-green, D_BMC1 dark-blue, BMC2 dark-purple, and D_BMC2 light-purple. (**C**) Daily average of seawater temperature profile during 2022. (**D**) Experimental design created by a licensed version of BioRender.

### Consortia composition

Two probiotic and two postbiotics assemblages were used in this experiment: Consortium-one (probiotic BMC1 and its postbiotic counterpart D_BMC1), and consortium-two (probiotic BMC2 and its postbiotic counterpart D_BMC2. The consortia-one consisted of a total of six bacterial strains, two *Pseudoalteromonas galatheae* and two *Cobetia sp.* strains isolated from *Pocillopora favosa*^91^; one *Halomonas sp.* strain isolated from *Stylophora pistillata*^92^, and one *Sutcliffiella sp.* strain isolated from *Galaxea fascicularis* (Linneaus, 1767), all collected in the CPV, Red Sea. The aforementioned BMC1 was previously used in an *in-situ* study by Delgadillo-Ordoñez and colleagues^55^ and the strains’ genomes were characterized by^93^. Their raw reads and assemblies are publicly available in the European Nucleotide Archive, under accession number PRJEB62849.

Consortium-two consisted of a total of four bacterial strains: one *Halomonas piezotolerans*, one *Pseudoalteromonas sp.*, and one *Pseudoalteromonas lipolytica,* isolated from *Acropora sp.*, and one *Cobetia sp.*, isolated from *P. favosa* from the CPV, Red Sea. The bacteria were selected based on beneficial traits (such as the presence of nifH, nirK and DMSP degradation genes, and the ability to produce urease, catalase and siderophores and to assimilate phosphate - as suggested by ^28^ from a Biovault of the Marine Microbiomes Laboratory, KAUST. Further information on host identification, positive beneficial traits, and strain identification is provided in Table S9.

The postbiotics D_BMC1 and D_BMC2 consist of their heat-killed probiotics counterparts. Their preparation is explained in the following session (Consortia preparation - Probiotics, postbiotic, and placebo).

### DNA extraction of BMCs’ strains, full-length 16S rRNA gene sequencing and strain identification

The BMCs strains’ genomic DNA was extracted using the Wizard® Genomic DNA purification kit (Promega Corporation, USA), following the protocol for Gram-positive and Gram-negative bacteria. The full-length 16*S* rRNA gene was PCR-amplified and sequenced using the 27F (5′-AGAGTTTGATCCTGGCTCAG-3′) and 1492R (5′-TACGGYTACCTTGTTACGACTT-3′)^94^ primers for taxonomic identification using the following cycles: one cycle of initial denaturation at 95°C for 5 min, 30 cycles of denaturation at 95°C for 1 min, annealing at 50°C for 1 min, extension at 72°C for 1 min, and one cycle of final extension at 72°C for 7 min. The PCR products were then Sanger-sequenced at the Bioscience Core Laboratory in KAUST, The forward and reverse sequences (1000–1500 bp) were processed to remove low-quality bases and generate contigs using the ChromasPro software. Ambiguities in the assembled sequences were resolved visually (either by choosing the base from the read with the cleaner signal or by changing the consensus base to “N”). Cleaned assembled DNA sequences from each of the BMC isolates were then identified using the EzBioCloud server^95^. The top-hit taxon, obtained from percent identity values, was used to estimate the taxonomy of the BMC isolates.

### Consortia preparation - Probiotics, postbiotic and placebo

The probiotic preparation was based on the strains’ growth curve, ensuring each strain was collected proportionally during their log phase according to their individual growth rates. Fresh bacterial cultures were collected according to their growth curve rates and washed three times with saline solution (3.5% NaCl). The cells of each strain were then re-suspended in 100 mL of saline solution for a final concentration 10^9^ cells/mL for the final probiotic consortium.

The postbiotics (here referred as dead_BMC consortia - D_BMC1 and D_BMC2) consisted of their respective probiotics heat-killed by autoclaving the final consortium for 20 minutes and plated in Marine Agar (HiMedia Laboratories, Mumbai, India) for checking its inactivation. The placebo consisted of a sterile 3.5% NaCl solution, the same solution that was used to wash and resuspend the bacterial cells for the other consortium preparations. A total of 30 mL of each consortium was aliquoted in Norm-Ject 50 mL Plastic Syringes for further inoculation. Probiotics and postbiotics consortia, as well as the placebo were prepared weekly, at a final concentration of ∼10^9^ cells/ml per probiotic and postbiotic treatment was and kept for up to 6 days at 4°C until inoculation, following the same methodology as previously described ^55^.

### Experimental design and sample collection

A total of 25 visually pale colonies within 7-9 m depth range, morphologically identified as *Acropora cf. valida* were tagged and distributed into five treatments, consisting of five colonies per treatment (Fig. 6B). The treatments were as follows: Placebo (colonies that were inoculated with a 3.5% NaCl sterile solution), BMC1 (colonies treated with the probiotic BMC1 consortium), D_BMC1 (colonies treated with BMC1 postbiotic consortium, i.e. heat-killed BMC1 consortium), BMC2 (colonies treated with the probiotic BMC2 consortium), and D_BMC2 (colonies treated with BMC2 postbiotic consortium, i.e. heat-killed BMC2 consortium) (Fig. 6D). An initial sampling time (T0) was conducted on September 15th, 2022, after the peak of temperature in the CPV, and a final sampling time (T1) 14 days after, on September 29th, 2022 (Fig. 1C). Probiotic and postbiotic inoculations took place in an intense manner for 4 consecutive days (starting two days after T0 sampling), followed by a 4 days interval and four more days of consecutive inoculations. Only after one day without any inoculation, the last sampling time (T1) was collected. A total of 30 mL of the respective treatments was applied over the colonies by using a 50 mL syringe.

Samples were collected by scuba divers wearing gloves and using pliers as previously described^96^. One fragment of ∼5-cm was collected for microbiome and metabolomics, as well as a fragment of ∼7-cm collected for photosynthetic efficiency (*Fv/Fm*) measurements from each colony. The fragments were placed into individual, sterile, opaque sampling bags (Whirl-Pak ®) and carried back to the boat. Right after the dive, fragments for microbiome and metabolomics analyses were transferred to individual 5 mL cryotubes, topped-up with DESS buffer (20% dimethyl sulfoxide - DMSO, 0.25 M ethylenediaminetetraacetic acid - EDTA, and saturated sodium chloride - NaCl, adjusted pH to 8.0) and then snap-frozen in liquid nitrogen. Microbiome/ metabolomic samples were then transferred to an ultrafreezer until further processing. The additional fragments collected for *Fv/Fm* measurements were maintained in their individual sampling bags (Whirl-Pak®), inside a dark cooler filled with local seawater until arrival at the Marine Microbiomes Laboratory, in KAUST, where their photosynthetic efficiency was accessed. The boat trip from the study site to the marina took approximately 30-40 minutes. Seawater temperature and salinity were monitored throughout the duration of the experiment using multiparameter CTDs (Ocean Seven 310 Multiparameter CTD, Idronaut). The daily minimum, maximum and mean seawater temperature (Fig. 6C) and mean salinity values are summarized in Table S1.

### Coral health assessment

Coral health status was assessed at the beginning of the experiment (T0) and at the end of the experiment (T1) by two different proxies: qualitative bleaching monitoring and the endosymbiotic algae photosynthetic parameters. For the qualitative bleaching monitoring, underwater pictures of the colonies were taken at T0 and T1 using a Sony Alpha 7 mk4. Pictures from the coral colonies were taken framing a Coral Health Chart to be used as a reference for coral color (University of Queensland)^97^. Four volunteers that have never seen the colonies’ pictures before performed a blind color scoring using the category “D” of the Coral Health Chart. It is worth mentioning that there was no reference to the different treatments in those pictures. The results of this assessment were then plotted in a heatmap for visualization.

Coral health was also assessed by the photosynthetic efficiency (*Fv/Fm*) of the endosymbiotic algae at the beginning (T0) and the end (T1) of the experiment. Samples collected to measure photosynthetic efficiency were dark adapted for at least 30 minutes after arriving to KAUST, and their *Fv/Fm* was measured using a Maxi Imaging-PAM *M-series* (Heinz Walz GmbH, Effeltrich, Germany), equipped with an IMAG-K4 camera and mounted with an IMAG-MAX/F filter. The measurements were conducted using the following settings: 6-7 light intensity, FM close to 0.14, Gain 2, Damping 2, Aperture 4-2.8 and Frequency 1. Factory standards were used for the remaining parameters. Measurements from three different points on each fragment were used to calculate the mean value. *Fv/Fm* changes over time, within the different treatments, were analyzed by a linear mixed effect model using the ‘*lmer*’ function from the “lme4” package on Rstudio (version 4.2.3)^98^. Replicates (n = 5) nested to treatment were treated as a random effect on the intercept to account for the non-independence of replicates with time. *Fv/Fm* was included in the model as a response variable, sampling time as a predictor variable, and treatment as a factor with five levels: Placebo, BMC1, D_BMC1, BMC2, D_BMC2. Pairwise contrasts were estimated by the “emmean” R package using the sampling times within treatment. One-way ANOVA tests were used to compare the different treatments against placebo. The package “ggplot” was used to generate the graphs. The data used as entry for the model and the script are available in the GitHub link in the data availability section.

### DNA extraction and bacterial 16S rRNA gene sequencing

Genomic DNA from all samples, from each sampling point (T0 and T1), was extracted. The DNeasy® Blood & Tissue kit (Qiagen) was used following their pre-treatment for Gram-positive bacteria, as follows: a 0.5 g coral fragment was placed directly into the lysis buffer and proteinase K was added. An overnight incubation was carried for approximately 16 hours, at 56°C, with a constant agitation of 650 rpm in a Thermomixer (ThermoFisher®). Subsequently, the manufacturer’s instructions were followed. A reaction with no biological material input was also performed to account for potential kit contaminants. DNA samples were stored at - 20°C until downstream analyses. DNA concentration and purity for all samples were quantified using the Qubit 2.0 Fluorometer High Sensitivity DNA Kit (Invitrogen™) and Nanodrop™ 8000 Spectrophotometer (Thermo Scientific™) and then normalized for 5 ng/mL.

To assess microbiome composition, samples had the hypervariable region V3-V4 of the 16*S* rRNA gene amplified by using the universal primers 341F 5’ CCTACGGGNGGC WGCAG 3’and 785R 5’ GAC TAC HVG GGT ATC TAA TCC 3’. Triplicate PCRs were performed for each sample (1 μl of input DNA) using Kapa HiFi HotStart Master Mix (Roche®), with a final primer concentration of 0.3 μM in a reaction with a final volume of 10 μl. Additionally, null template PCRs (no template DNA input) and a negative control from the extraction step were run to account for putative kit contaminants. The thermal cycling conditions were as follows: initial denaturation at 95 °C for 3 minutes, 35 cycles of 95°C for 30 secondsg, 62 °C for 30 seconds, 72 °C for 30 seconds, followed by a final extension at 72 °C for 5 minutes. Amplification was confirmed by running the 96 well plates into the QIAxcel Advanced System (QiaGen) that performs automated capillary electrophoresis to determine PCR quality and integrity. Triplicate PCRs from each sample were pooled and samples cleaned using AMPure XP beads (Beckman Coulter) to purify the amplicon from primer dimers and free primers. Samples were then indexed using Illumina sequencing adapters (Nextera XT) dual index primers to encode each sample, libraries were purified using the AMPure XP reagent (Beckman Coulter™) according to Illumina specifications, and the excess primers, nucleotides, salts, and enzymes were removed using the washing procedure. Libraries were screened before sequencing, in which each library was pooled based on Quant-iT™ dsDNA High-Sensitivity Assay calculated molarity. The pool quality was assessed by Agilent TapeStation electrophoresis System to determine the libraries concentrations and average size. In addition, a qPCR-based quantification to find the DNA target and to derive a quantitative estimate of the initial template concentration for the pool was conducted, to achieve the sequencing performance the control/assurance (QC/QA) systems recommend for NGS. Libraries were then sequenced at the KAUST Bioscience Core Laboratory using the NovaSeq 6000 platform (Illumina) and 250 bp paired-end reads were generated.

### Bacterial community analysis

The *DADA2* pipeline^99^ was used to infer Amplicon Sequence variants (ASVs) from the 16*S* rRNA gene-based amplicon read-libraries. Briefly, the raw reads were decontaminated of phiX and adapter-trimmed using the BBDuk tool from the BBMap suite (Bushnell B, http://sourceforge.net/projects/bbmap/). The PCR primers were removed using the cutadapt tool^100^. After merging the forward and reverse reads via the mergePairs function of DADA2, the sequences were analyzed under the pseudo-pooling mode by following the DADA2 (version 1.22) workflow and by using the SILVA database, version 138.1^101^. All potential contaminant ASVs that were identified by the decontam tool^102^ were removed from the analysis. ASV raw counts are available on the Zenodo repository (as indicated in the data availability section).

The microbial ASVs counts generated by DADA2 were imported into phyloseq version 1.42.0^103^ on RStudio (version 2021.09.0) for downstream analyses. First, ASVs corresponding to mitochondria and chloroplast were removed, followed by the depletion of singleton ASVs using the phyloseq function ‘*prune_taxa’*, resulting in 10,688 ASVs. Rarefied samples were used for calculating alpha diversity analysis only, in which samples were rarefied using 300,000 reads as a cut-off based on the plateau observed on the ASV rarefaction curve (Fig. S5), in which 9 samples were removed because they presented lower sequencing depth than the determined cut-off: F3 (T0 PT_BMC1_3), F8 (T0 BMC1_3), F14 (T0_D_BMC1_4), F24 (T0 D_BMC2_4), F25 (T0_D_BMC2_5), F32 (T1 PT_BMC1_2), F37 (T1_BMC1_2), F48 (T1 BMC2_3), F54 (T1_D_BMC2_4). A total of 5,157 ASVs were retained after read-rarefaction. Alpha diversity was estimated with the function ‘*estimate_richness*’ on *phyloseq*. Statistical comparisons of alpha diversity-indices between sampling times within treatments and between treatments at T1 were performed using Wilcoxon tests, since Shapiro-Wilk test indicated non-normality for data distribution (p-value < 0.05). The non-rarefied ASV dataset was used with ‘betadisper’ command from the ‘vegan’ package to check for differences in the homogeneity of variances^104^ Non-metric MultiDimensional Scaling (nMDS) were used to visualize patterns in beta diversity, and permutational multivariate analyses of variance (PERMANOVA) were applied to Bray-curtis matrixes to test for differences in the bacterial communities between treatments in each sampling time. For all the statistical analysis, p-values were adjusted to account for False Discovery Rate

(FDR) for multiple comparisons, using the Benjamini and Hochberg (BH) correction^105^. To further explore bacterial differential abundance between treatments in T1 we used the function DESeqfrom the package DESeq2^106^ in R, on a model of negative binomial distribution; a Wald test with parametric fitting of dispersions to the mean intensity was used to estimate differentially abundant taxa, using a cutoff value of p = 0.01. Additionally, the ASV sequences from all treatments from T1 were queried for the presence of the 16*S* rRNA gene sequences from each of the BMCs strains, using BLASTn^107^. From the non-rarefied ASV dataset we identified 115 ASVs that shared ≥99% sequence identity (at >400 bp query length) with the 16*S* rRNA genes of the BMC strains. This set of ASVs was then used to evaluate enrichment between sampling times and treatment using Kruskall-Wallis followed by Dunn test. Importantly, no ASVs with ≥99% sequence identity to *Pseudoalteromonas lipolytica* were detected, and therefore this strain was not included in the plots of this analysis.

### Metabolomics

Approximately 500 mg of each coral sample was used to extract the metabolites. Initially, the fragments were homogenized with 80% methanol (1 mL) in a bead blaster and subjected to three cycles of 1 min 30 sec, at 4°C and 3,000 rpm. The resulting extract underwent further processing, being placed in a Thermomixer for 1.5 hours at 4°C and 1,400 rpm, sonicated at 4°C for 20 minutes, and subsequently centrifuged at 15,000 rpm for 10 minutes at 4°C. The supernatant obtained was then collected and lyophilized. Following the drying step, the powder was resuspended in a 400 µL Deuterium oxide solution containing a 20 mM potassium phosphate and Trimethylsilylpropanoic acid standard (TSP) mixture to stabilize the pH (7.4) and allow for comparison between samples. For NMR data acquisition, 200 µL of sample was added to 3-mm NMR tubes and then added to a Bruker 700 MHz ADVANCE NEO NMR spectrometer equipped with a Bruker TXO (^1^H/^13^C/^15^N/^2^H) cryogenic probe (BrukerBioSpin, Rheinstetten, Germany) at 298K. Transformed spectra underwent processing for phase correction using Topspin 4.1.4 (Bruker BioSpin) and calibrated to the proton signal of TSP at 0.00 ppm. Following this, baseline correction was carried out on Chenomx profiler (version 9.2). The analysis of metabolite molecules involved annotating their profiles using standard peaks from Chenomx. Additionally, 2-D NMR spectra was used to confirm identities for the annotation of metabolites and confirmed with literature^108^. Further, all comparative concentrations of metabolites were batch-exported and imported into both R and Metaboanalyst software for comprehensive data exploration and statistical analysis. To ensure the comparability of individual samples, normalization was carried out using the weight values collected prior to extraction, then they underwent log-transformation and pareto scaling^109^. The differences between treatments were plotted by using Principal Component Analysis (PCA) to visualize variations. Overall multivariate differences among treatments were tested using PERMANOVA on Euclidean distances with 999 permutations. For each metabolite (univariate analyses), we first assessed assumptions by fitting a one-way model (individual_metabolite_value ∼ treatment) and testing normality of ANOVA residuals with the Shapiro–Wilk test and homogeneity of variance with Levene’s test. If tests were non-significant, a one-way ANOVA was used and pairwise differences identified Tukey’s HSD. If normality assumption failed, Kruskal-Wallis test was used and pairwise contrasts evaluated with Dunn’s test using BH adjustment. To control for false discovery rate across metabolites, primary test p-values (ANOVA or Kruskal) were adjusted using BH. Analysis in R were conducted using vegan, car, FSA and tidyverse packages.

To access positive and negative correlations between the presence of the probiotics or postbiotics’ strains and the metabolites generated by the NMR analysis, the data was first transformed using centered log ratio (CLR) transformation^110^, then submitted to Spearman’s rank correlation coefficient (significance thresholds were p<0.05) using the ‘corrplot’ R package ^111,112^. The genera from the strains composing the consortia-one and -two (*Pseudoalteromonas*, *Cobetia* and *Halomonas*) were filtered from ASVs counts and used to imply correlations with the normalized metabolites matrix within treatments (files available in Zenodo repository). ASVs classified as *Sutcliffiella* were not detected in the dataset. Since *Sutcliffiella* was previously classified as *Bacillus*, we attempted to check for correlations between *Bacillus* instead. However, as no significant correlations were found between *Bacillus* and any of the detected metabolites, neither S*utcliffiella* or *Bacillus* was not further reported in the main correlation plots.

### Statistics and reproducibility

All statistical analyses were performed using R (v4.3.1). The statistical tests used for each analysis are described in its respective methods sections. All analyses were conducted with 5 biological replicates per treatment group, as specified in each figure legend. Microbiome analyses were performed using the Phyloseq, vegan, and DESeq2 packages. Bray–Curtis distance matrix was used for beta diversity, and PERMANOVA (adonis2, 999 permutations) was applied to test for group differences. Differential abundance was assessed using DESeq2, of which p-values were adjusted for multiple comparisons using the Benjamini–Hochberg method, with significance defined as FDR <0.01. All raw sequence reads from the study were deposited in the European Nucleotide Archive (ENA) under the study accession number PRJEB89680. The analysis scripts are available at Zenodo (10.5281/zenodo.17116572).

## Acknowledgements

The authors would like to thank the KAUST Coastal and Marine Resources Core Lab (CMR) staff for help with field work logistics and support. Carolina Bocanegra is thanked for assistance with library preparation. We are grateful to the staff of the KAUST Bioscience Core Laboratory, especially Rubén Díaz-Rúa, for troubleshooting the molecular work. Francesca Benzonni is thanked for helping with the taxonomic identification of the coral colonies. We also thank the volunteers who helped with performing the blinded bleaching scores.

## Author contribution

EPS and RSP conceived the study. EPS, IR and NGB performed fieldwork and collected data. JS and ASR supported the culturing and production of the probiotics/postbiotics. EPS did the molecular work. EPS, FG, CPA, NGB, JC, IR and RSP analyzed and/or interpreted the microbiome data. EPS, IR, LB, US and LJ were involved in processing and/or analyzing the metabolomics. ND-O performed the photogrammetry analysis. ASR, RSP, LJ and MJ provided financial support. EPS and RSP wrote the initial draft. All the authors made contributions to the final manuscript, read and accepted previous to submission.

## Data availability

All data needed to evaluate the conclusions in the paper are present in the paper and/or the Supplementary Materials. All raw sequence reads from the study were deposited in the European Nucleotide Archive (ENA) under the study accession number PRJEB89680. Scripts and additional material used to analyze the data are available on Zenodo (10.5281/zenodo.17116572).

## Supplementary figures

**Fig. S1:**
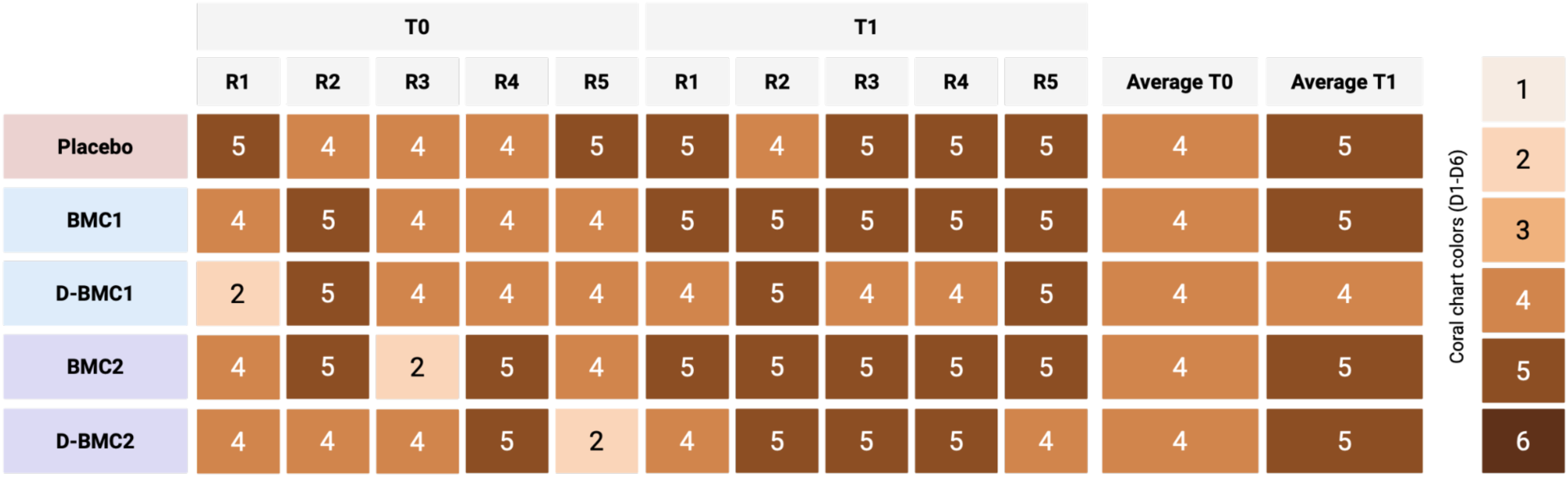
Color score of the colonies used in each of the treatments by time point.

**Fig. S2:**
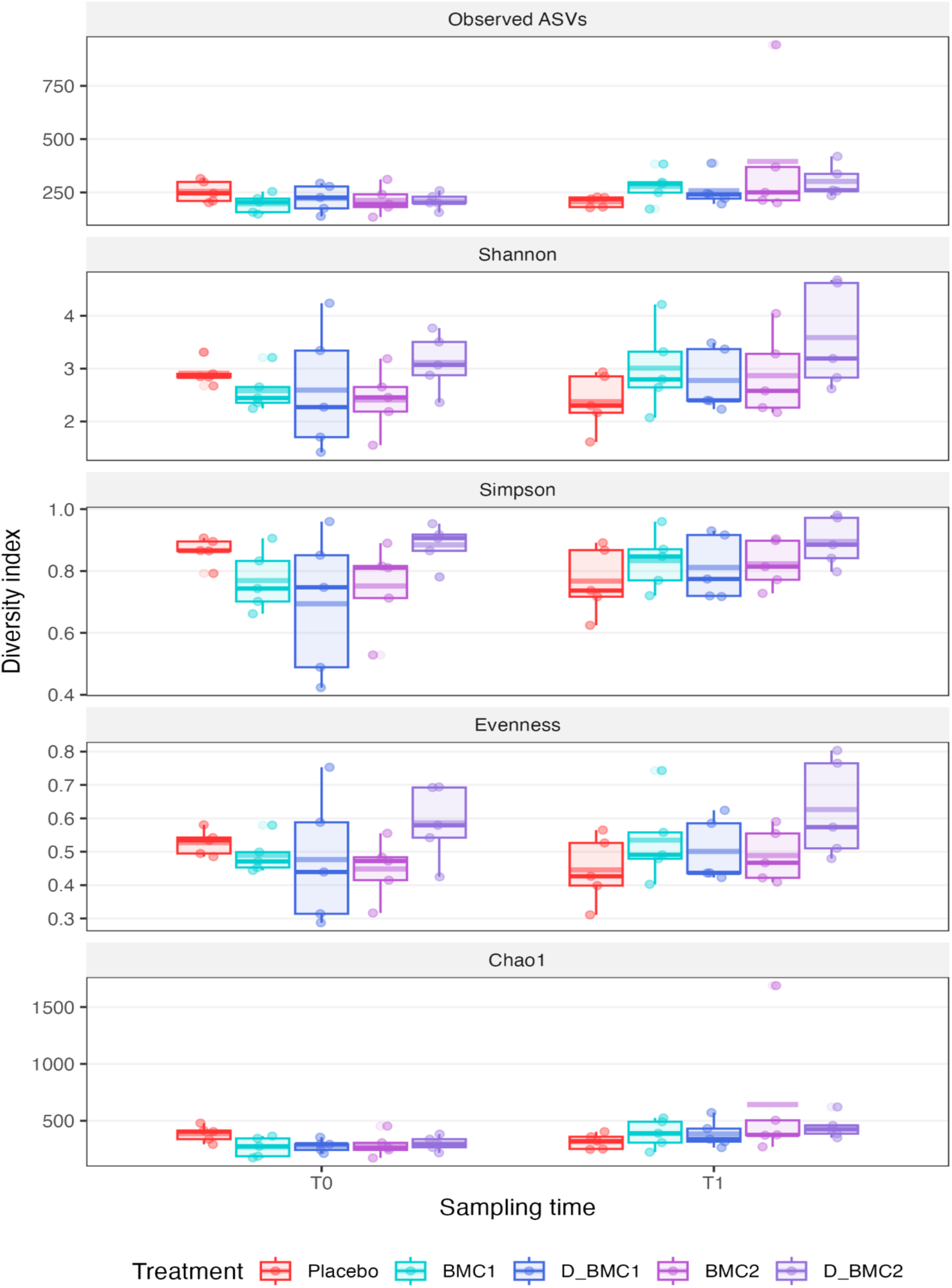
Alpha diversity metrics across treatments and sampling times. Pairwise comparisons between categories were calculated employing Wilcoxon tests. Categories were considered significantly different at p ≤ 0.05.

**Fig. S3:**
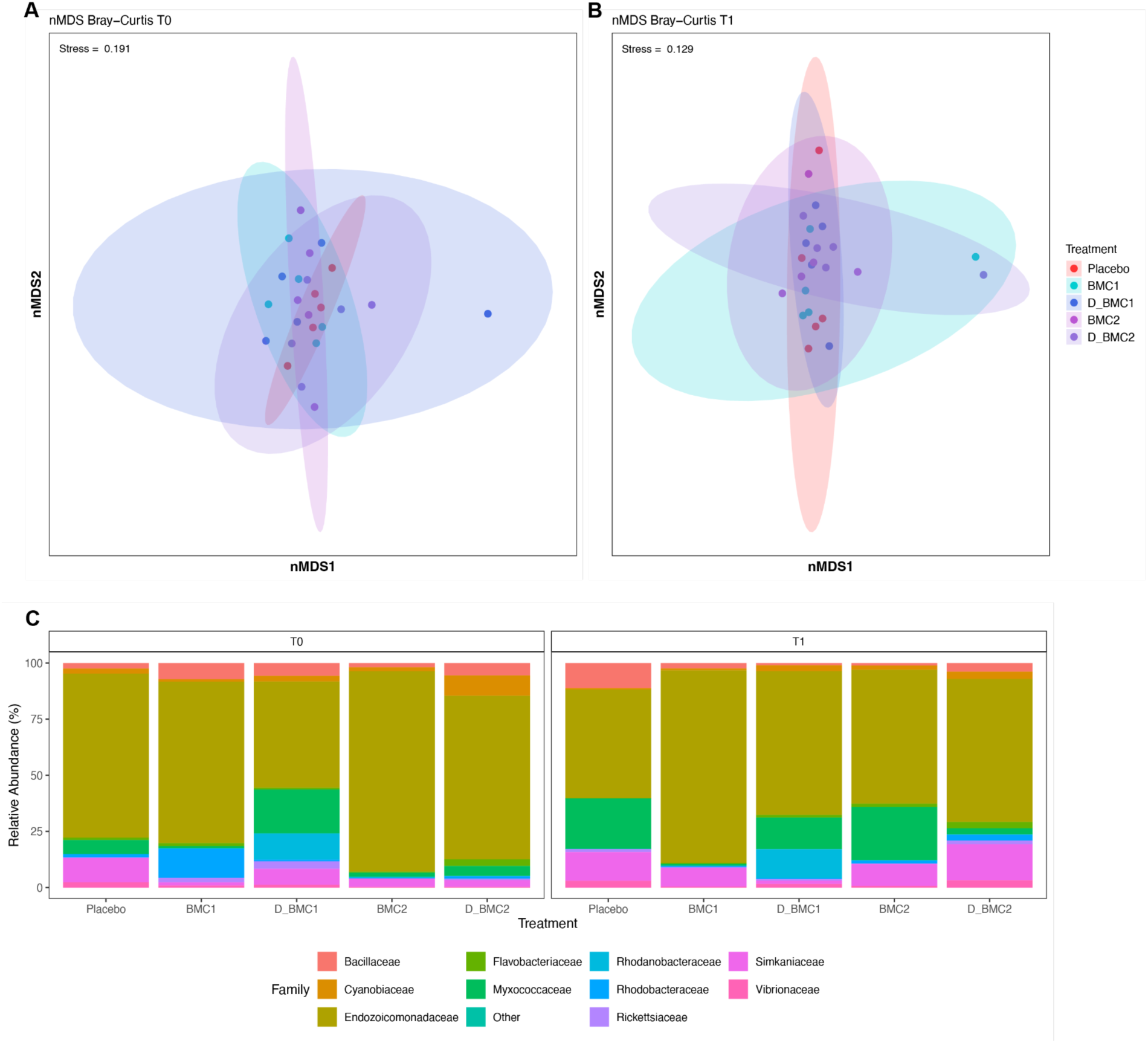
(**A**) Sample distribution on Nonmetric multidimensional scaling ordination (nMDS) using Bray-Curtis distance matrices by treatment in T0 and (**B**) T1. (**C**) Barplots of bacterial families distribution identified by their 16S rRNA gene sequencing by treatment and sampling time.

**Fig. S4:**
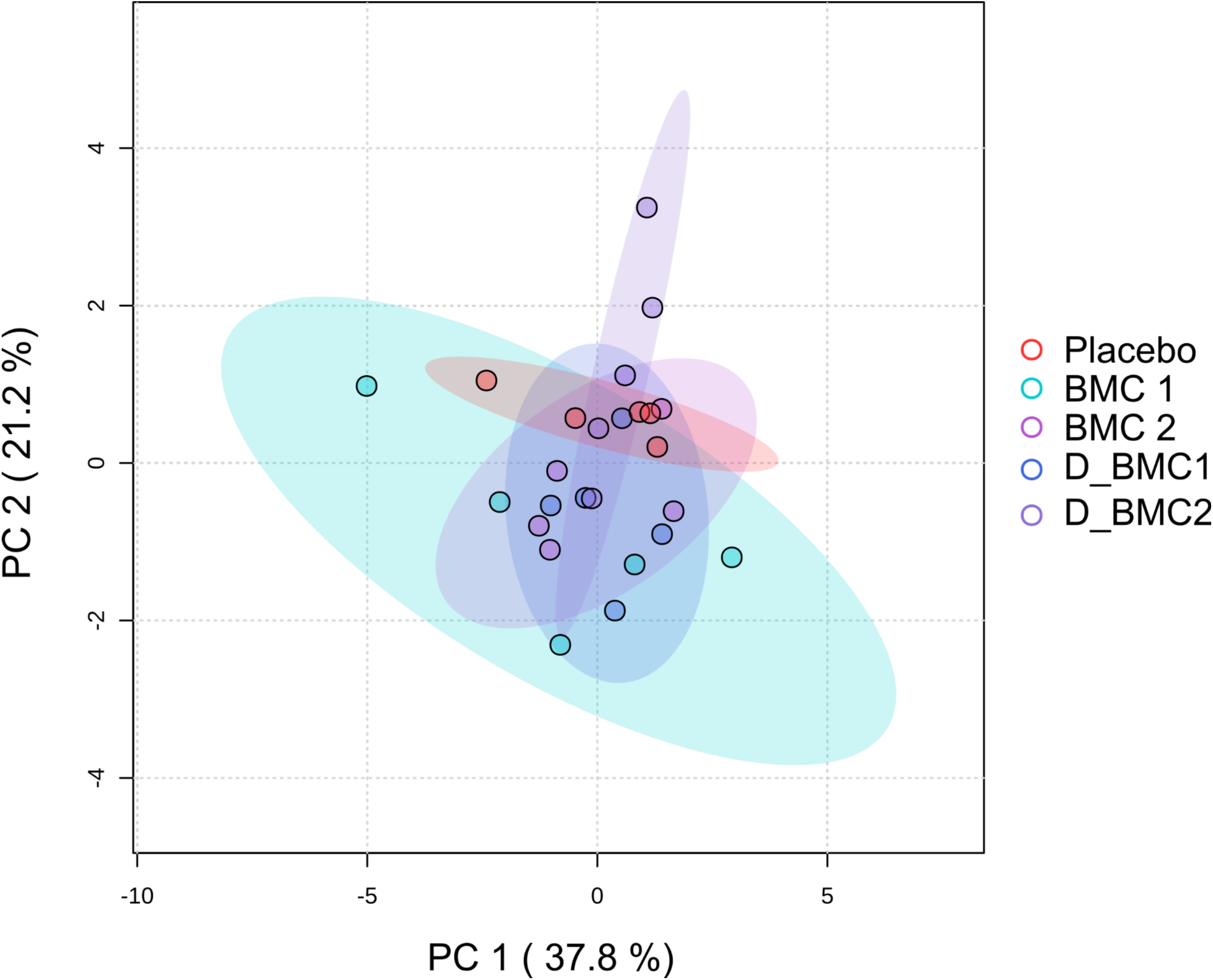
Ordination plots of PCA from data of ^1^H NMR spectra before the use of microbial treatments (T0).

**Fig. S5:**
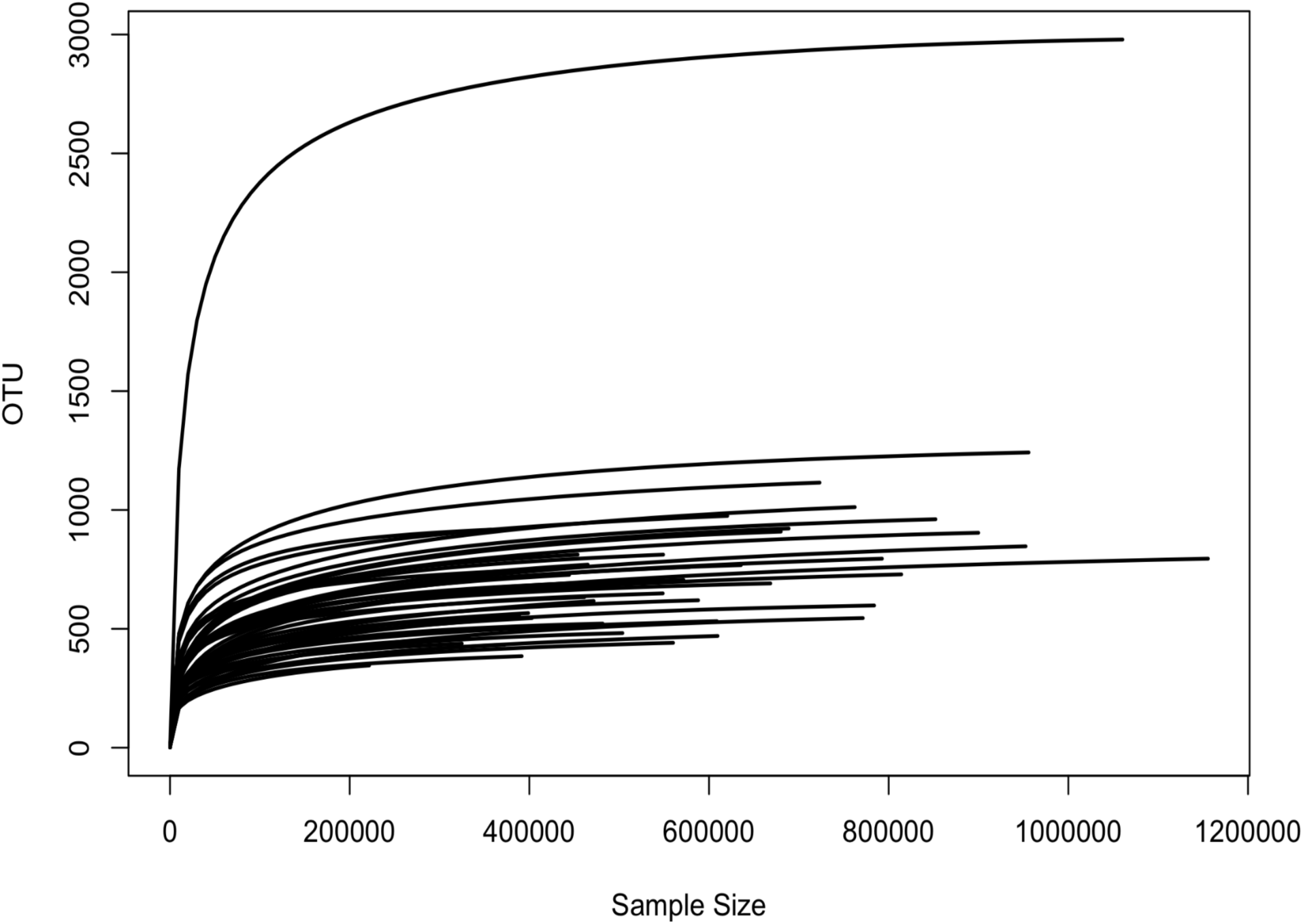
ASV rarefaction curve from the 16*S* rRNA gene sequencing data.

## Supplementary Tables

**Table S1:** Daily minimum, maximum and mean values of seawater temperature, and mean salinity values of the study site during the period of the experiment.

| Sampling Time | Date | Daily Average Temperature (°C) | Daily Minimum Temperature (°C) | Daily Maximum Temperature (°C) | Average Salinity (PSU) |
| --- | --- | --- | --- | --- | --- |
| T0 | 15/09/2022 | 31.95 | 31.75 | 32.19 | 39.44 |
|  | 16/09/2022 | 31.54 | 31.32 | 31.80 | 39.20 |
|  | 17/09/2022 | 31.67 | 31.37 | 32.13 | 39.30 |
|  | 18/09/2022 | 31.50 | 31.25 | 31.81 | 39.38 |
|  | 19/09/2022 | 31.39 | 31.13 | 31.77 | 39.35 |
|  | 20/09/2022 | 31.24 | 30.91 | 31.71 | 39.34 |
|  | 21/09/2022 | 30.99 | 30.67 | 31.34 | 39.32 |
|  | 22/09/2022 | 30.98 | 30.70 | 31.36 | 39.27 |
|  | 23/09/2022 | 30.87 | 30.49 | 31.32 | 39.41 |
|  | 24/09/2022 | 30.87 | 30.57 | 31.16 | 39.47 |
|  | 25/09/2022 | 31.05 | 30.79 | 31.33 | 39.08 |
|  | 26/09/2022 | 31.14 | 30.91 | 31.38 | 39.23 |
|  | 27/09/2022 | 31.16 | 30.87 | 31.45 | 39.31 |
|  | 28/09/2022 | 31.13 | 30.89 | 31.40 | 39.29 |
|  | 29/09/2022 | 31.21 | 30.97 | 31.46 | 39.49 |
| T1 | 30/09/2022 | 31.24 | 30.99 | 31.52 | 39.50 |

**Table S2:** Pairwise comparison of Fv/Fm between sampling times per treatment. Degrees of freedom method used was Kenward-Roger. Sampling time was selected as a contrast variable and the different treatment included as a factor with five levels was included as a treatfact. Parameter estimated (estimate), standard error (SE), degrees of freedom (df), t ratio (t ) and p-value (p) are provided.

| contrast | treatfac | estimate | SE | df | t | p |
| --- | --- | --- | --- | --- | --- | --- |
| T0 - T1 | Placebo | 0.05 | 0.03 | 19.79 | 2.12 | 0.047* |
| T0 - T1 | BMC1 | -0.00 | 0.03 | 19.79 | -0.02 | 0.985 |
| T0 - T1 | D_BMC1 | 0.07 | 0.03 | 19.79 | 2.71 | 0.014* |
| T0 - T1 | BMC2 | 0.01 | 0.02 | 17.65 | 0.57 | 0.578 |
| T0 - T1 | D_BMC2 | 0.03 | 0.02 | 17.65 | 1.31 | 0.208 |

**Table S3:** Distribution of ASVs with ≥99% identity with BMCs strains in the different treatments and sampling time (mean and median). The table also provides the results of statistical analysis (Kruskall Wallis) of each bacterial strain by time among treatments followed by Dunn test for identifying enrichment of BMCs strains in different treatments at T1. (Excel file)

**Table S4:** Kruskal-Wallis statistical test results of alpha diversity (Shannon, Observed ASV, Simpson, Chaos and Evenness indexes) of 16S rRNA ASVs in each sampling time. (Excel file)

**Table S5:** Results of PERMANOVA test based on Bray-Curtis dissimilarities of 16S rRNA ASVs.

| Sampling time T0 | Df | SumOfSqs | R2 | F | Pr(>F) | Test |
| --- | --- | --- | --- | --- | --- | --- |
| Model | 4 | 1.75092877 | 0.18059391 | 1.10198052 | 0.219 | PERMANOVA |
| Residual | 20 | 7.94446338 | 0.81940609 | NA | NA | PERMANOVA |
| Total | 24 | 9.69539215 | 1 | NA | NA | PERMANOVA |
| Sampling time T1 | Df | SumOfSqs | R2 | F | Pr(>F) | Test |
| Model | 4 | 1.65952572 | 0.17627295 | 1.06997185 | 0.29 | PERMANOVA |
| Residual | 20 | 7.75499708 | 0.82372705 | NA | NA | PERMANOVA |
| Total | 24 | 9.4145228 | 1 | NA | NA | PERMANOVA |

**Table S6:** Relative abundance of bacteria families in the different sampling times and treatments. (Excel file)

**Table S7:** DESeq results and statistics, including individual FDR adjusted p-values, log2FolderChange (LFC) of the differentially abundant taxa per treatment at T1. (Excel file)

**Table S8:** Statistical tests based on the metabolites detected by 1H NMR analysis. PERMONOVA was used for detecting differences in the metabolic profile between treatments. ANOVA or Krukall-Wallis (depending on distribution normality) was used to determine differences in the concentration of individual metabolites among treatments, which was followed by pairwise comparison between treatments by Tukey or Dunn’s test with padjusted. (Excel file)

**Table S9:**
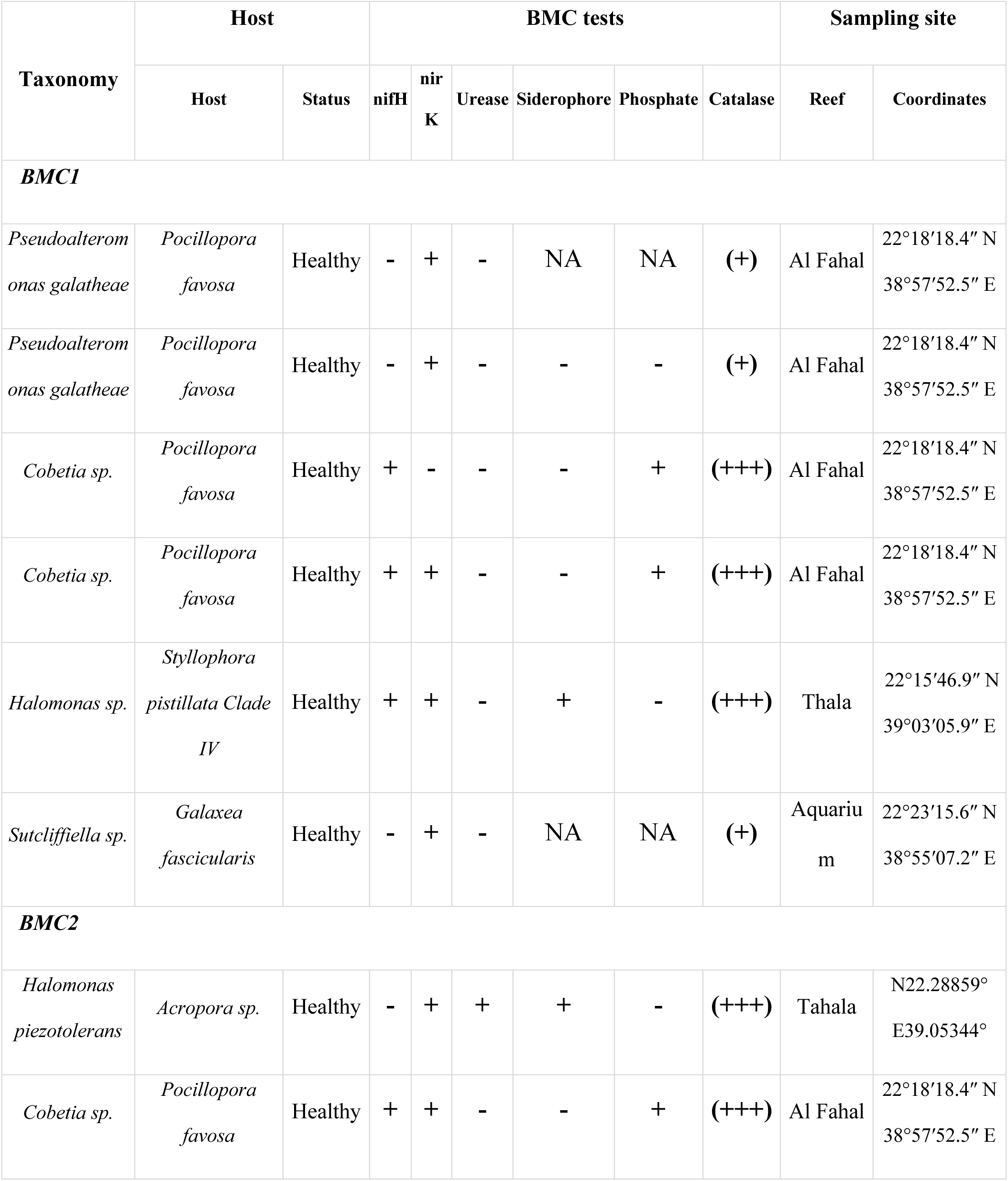
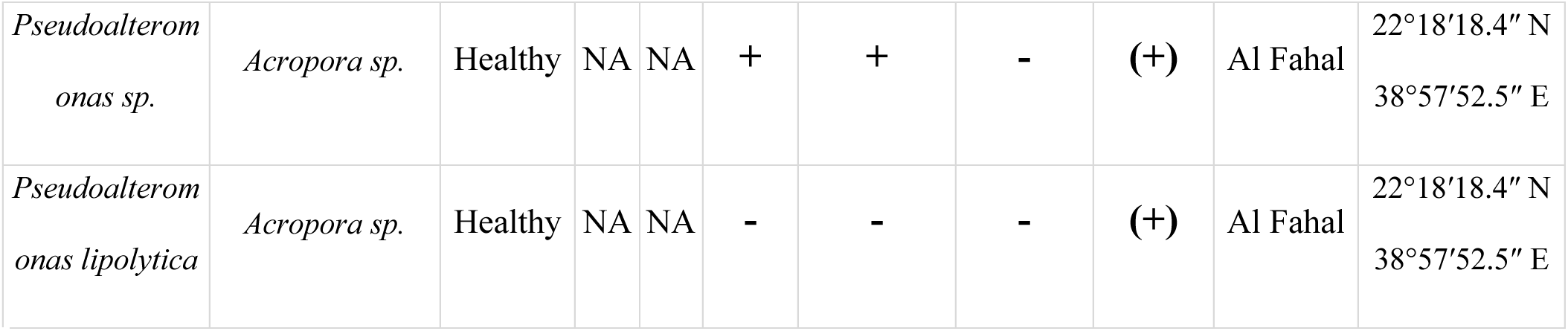
Information on the bacteria strains used as probiotics in both BMC1 and BMC2.

